# Bimodality in pan-cancer proteomics reveals new opportunities for biomarker discovery

**DOI:** 10.1101/2025.03.03.641218

**Authors:** Wen Jiang, Damian Bikiel, Jan Zaucha, Jixin Wang, Sanddhya Jayabalan, Yongjin Li, Matthew Sung, James Harper, Jimmy Zhao, Krishna Bulusu, Wenyan Zhong

## Abstract

Bimodal protein expression, characterized by the distribution of protein expression with two modes, is linked to phenotypic variation across various biological systems. Whereas previous studies focused on RNA expression data, we developed a bimodality model tailored for proteomics to enhance the identification of cancer-associated biomarkers and targets, facilitating precision oncology. We analyzed proteomics data from various cancer types and identified 2401 tumor-associated bimodal proteins. These proteins were evaluated for pathway enrichment, revealing significant associations with critical cancer pathways, such as metabolism of non-essential amino acids, interaction between the extracellular matrix and its receptors on the cell surface, and central carbon metabolism in cancer. Utilizing an AI-enhanced knowledge graph, we further delineated common patterns among pan-cancer tumor-associated bimodal proteins. A case study on the bimodal expression of TROP2 in colon adenocarcinoma highlighted upregulation of MYC and WNT/β-catenin signaling pathways and down-regulation of inflammatory and interferon-related pathways in the TROP2-high group. The biological difference between TROP2-high and TROP2-low groups underscored its significance in determining cancer heterogeneity and differences in cancer vulnerability, which can inform treatment decisions. Our findings show the value of proteomics in uncovering novel biomarkers and advancing precision medicine, setting a precedent for further multi-omics integration and clinical validation.

## Introduction

Identifying cancer-associated targets and biomarkers from high-throughput genomics, transcriptomics, and proteomics is a fundamental process in precision oncology.^1–6^ The established method for detecting cancer-associated targets, biomarkers, and oncogenes as well as tumor suppressor genes is through the comparison of tumor samples with matched healthy tissues.^7–10^ Recently, however, the strategy of recognizing the activation of distinct gene programs in phenotypically diverse tumor tissues, which is reflected as two distinct peaks in RNA sequencing (RNA-seq) expression profiles, has emerged as an alternative method.^11–13^ One advantage of utilizing a bimodal gene as a biomarker lies in the ability to effectively categorize patients into two clearly distinct expression states. This feature enhances the ease of interpretation, reproducibility, and application of the biomarker in clinical settings. An example is estrogen receptor ESR1, whose bimodal expression classifies breast cancer patients into two subtypes (ER positive and negative) and leads to different treatment strategies.^14,15^

Previous studies leveraging different bimodality detection methods, such as model-based clustering,^13,16^ COPA,^9^ and SIBER,^17^ on RNA-seq data sets have unveiled the widespread occurrence of bimodal gene expression across various biological contexts.^18^ This dichotomous expression phenomenon plays a pivotal role in developmental biology, stem cell pluripotency, and the emergence of cellular diversity within tissues.^16,19^ Moreover, the bimodal distribution of gene expression levels has been implicated in disease pathogenesis, including cancer, where it may influence tumor heterogeneity, drug resistance, and metastatic potential.^13,20–24^

Deciphering the mechanisms underlying bimodal gene expression offers promising avenues for therapeutic intervention and precision medicine.

However, there are two major limitations in previous studies. First, despite the identification of significant cancer genes with bimodal expression patterns, most studies utilize RNA expression measured by RNA-seq.^13,17,25,26^ As reported in the literature, correlations between RNA and protein can be as low as 40% due to multiple factors, including protein translation efficiency and stability, mRNA transcription rate, and the biochemical diversity of proteins.^27,28^ In contrast to RNA-seq, proteomics can provide quantification of proteins that serve as the main executors of biological functions in cells and directly influence cellular processes and disease phenotype. A second challenge in previous studies is the difficulty of distinguishing between true bimodal expression and artifacts resulting from technical noise or insufficient sample sizes.^16,17^ These limitations underscore the need to develop an appropriate bimodality model that can reliably identify bimodal expression patterns in proteomics data sets.

Recent advancements in proteogenomics have provided unique opportunities for understanding the molecular intricacies of cancer biology.^3,10,29–40^ The National Cancer Institute’s (NCI’s) Clinical Proteomic Tumor Analysis Consortium (CPTAC) has profiled large collections of clinical tumor samples and normal adjacent tissues from cancer patients,^3,10,29–40^ providing high-quality mass spectrometry–based proteomics data. This represents an ideal resource for identifying bimodal protein expression patterns. In this study, we constructed a bimodality model to identify bimodal protein expressions in CPTAC pan-cancer proteomics data^41^ and identified 2401 tumor-associated bimodal proteins (TABPs) (Figure 1). To further understand the disease relevance of these novel TABPs, we leveraged an AI-boosted knowledge graph^42^ to identify shared biological functions of 16 pan-cancer TABPs. As an example, we performed a detailed case study on TROP2, a significant TABP identified in colon adenocarcinoma,^10^ to uncover its potential mechanisms and disease implications. Our findings highlight the unique disease biology associated with bimodal TROP2 protein expression and demonstrate its potential as a biomarker to inform therapeutic interventions.

**Figure 1.**
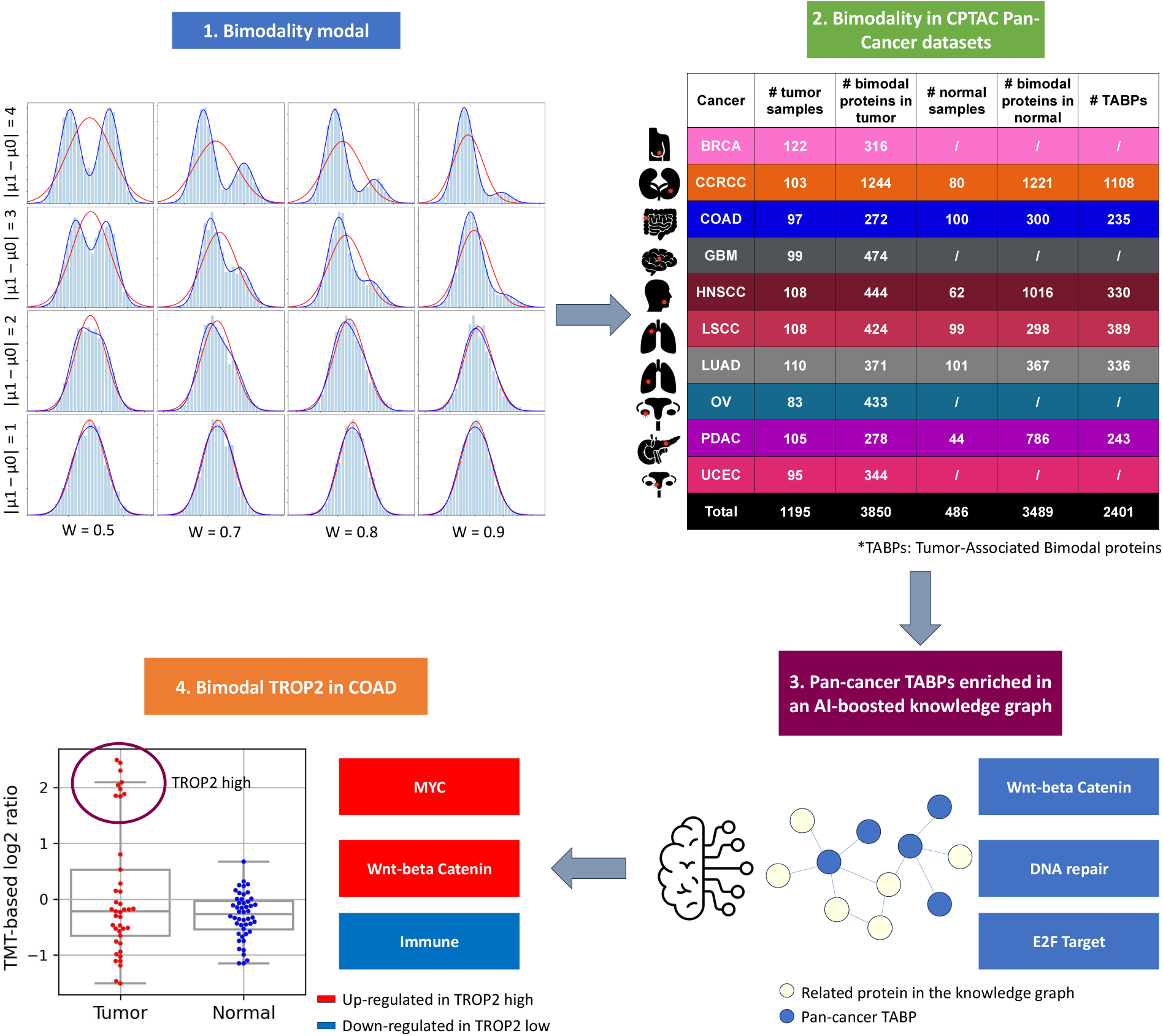
Workflow to (1) construct a bimodality model, (2) identify bimodal protein expressions in CPTAC pan-cancer proteomics data comprising 10 tumor cohorts and 6 normal cohorts, (3) capture the common patterns of pan-cancer tumor-specific bimodal proteins using an AI-based knowledge graph, and (4) explore biology behind TROP2 bimodality in the COAD cohort.

## Results

### Development of a bimodal model for proteomics data

To comprehensively understand the scope of bimodal protein expressions, we implemented a bimodal model for proteomics data (Figure 1). We first calculated three different bimodality metrics, (1) the bimodality index (*BI*), (2) the difference in Bayesian information criteria (*BIC diff*), and (3) kurtosis (*kurt*), on the simulation data sets (sample number *N* = 3000 or 100) with different combinations of mean differences and weights (Methods; Supplementary Figure S1).

Supplementary Figure 1A demonstrates a set of simulation data sets showing that, with large mean differences relative to variances and a sufficient sample size (*N* = 3000), bimodal distributions are easily identifiable. In such cases, *BI* exceeded 1, *BIC diff* ranged from 0 to – 1100, and *kurt* varied between –1.3 and 0. In contrast, indistinguishable cases featured *BI* < 1, *BIC diff* 0–50, and *kurt* approximately 0. Increasing separation in the first column led to rising *BI* and decreasing *BIC diff* and *kurt*. We observed that positive *kurt* typically indicated outliers, whereas negative *kurt* suggested evenly distributed mass clusters. Given that imbalanced bimodality, which is common in real-world data, often resulted in *kurt* near 0, we excluded *kurt* from our bimodality model to avoid misinterpretations.

A major challenge in bimodality identification is limited sample sizes, which may be caused by experimental design or missing values. Supplementary Figure S1B illustrates this challenge with a set of simulation data sets with only 100 data points (*N* = 100). Reduction of sampling from 3000 to 100 made it more difficult to discern cases previously categorized as easy. Despite smaller sample sizes, *BI* remained above 1 and *BIC diff* ranged from –38 to –16, indicating that these metrics can still effectively identify bimodal distributions under constrained conditions.

Next, we systematically evaluated *BI* and *BIC diff* using a benchmarking data set of 100 known bimodal and unimodal distributions (50:50; Methods, Supplementary Figure S2). We sampled different sizes from the benchmarking data set (sample number *N* = 1000, 300, 100, 70, 50, 30) and separately computed the true-positive rate (TPR) and false-positive rate (FPR) at different thresholds for *BI* and *BIC diff* (Supplementary Table S1, Supplementary Figure S3). When we used the max(TPR – FPR) as the cutoff, *BIC diff* showed slightly higher TPRs and lower FPRs than *BI* at all sample sizes. To enforce stricter standards on FPRs, we decided to use both *BI* and *BIC diff* metrics and apply different thresholds according to the actual number of samples with protein quantifications in the proteomics data sets (Table 1).

**Table 1.** Bimodality thresholds for proteomics data sets with different sample sizes.

| ***N*** | ***BIC diff* threshold** | ***BI* threshold** |
| --- | --- | --- |
| 30–50 | $\leq 3$ | $\geq 1.35$ |
| 50–100 | $\leq 4$ | $\geq 1.25$ |
| $\geq 100$ | $\leq 5$ | $\geq 1.1$ |

### Prevalence of bimodality in CPTAC pan-cancer proteomics data

From the large-scale proteomics data of the CPTAC pan-cancer data sets, we analyzed 10 tumor cohorts: breast cancer (BRCA),^3^ clear-cell renal cell carcinoma (ccRCC),^30^ colon and rectal cancer (COAD),^43^ glioblastoma (GBM),^38^ head and neck squamous-cell carcinoma (HNSCC),^44^ lung squamous-cell carcinoma (LSCC),^45^ lung adenocarcinoma (LUAD),^46^ ovarian cancer (OV),^39^ pancreatic ductal adenocarcinoma (PDAC),^29^ and uterine corpus endometrial carcinoma (UCEC).^47^ We identified bimodal proteins in the 10 tumor cohorts as well as in the 6 matched normal cohorts (ccRCC, COAD, HNSCC, LUAD, LSCC, and PDAC) using our bimodality model (Methods; Figure 1, Supplementary Table S1).

To focus our study on tumor-associated bimodality, we defined TABPs as bimodal proteins that are found in the tumors but not present in the matched normal tissues (2401 of 15,098 proteins; Methods, Supplementary Table S1). Pathway enrichment analysis revealed a significant association of TABPs with arginine and proline metabolism, extracellular matrix (ECM)-receptor interaction, central carbon metabolism in cancer, focal adhesion, and protein processing in endoplasmic reticulum in different cancer types (Figure 2A, Supplementary Table S2). For example, 15 ccRCC-specific TABPs (CD47, COL6A3, FN1, HSPG2, ITGA11, ITGAV, ITGB4, ITGB7, LAMA2, LAMB3, LAMC2, NPNT, THBS1, VTN, VWF; Supplementary Figure S4A) overlapped with the ECM-receptor interaction gene set, which primarily consists of various fibrillar collagens, fibronectin, and basement membrane components like laminin, collagen IV, and heparan sulfate proteoglycan.^48^ The bidirectional interaction between cells and the ECM influences cell adhesion and migration.^49^

**Figure 2.**
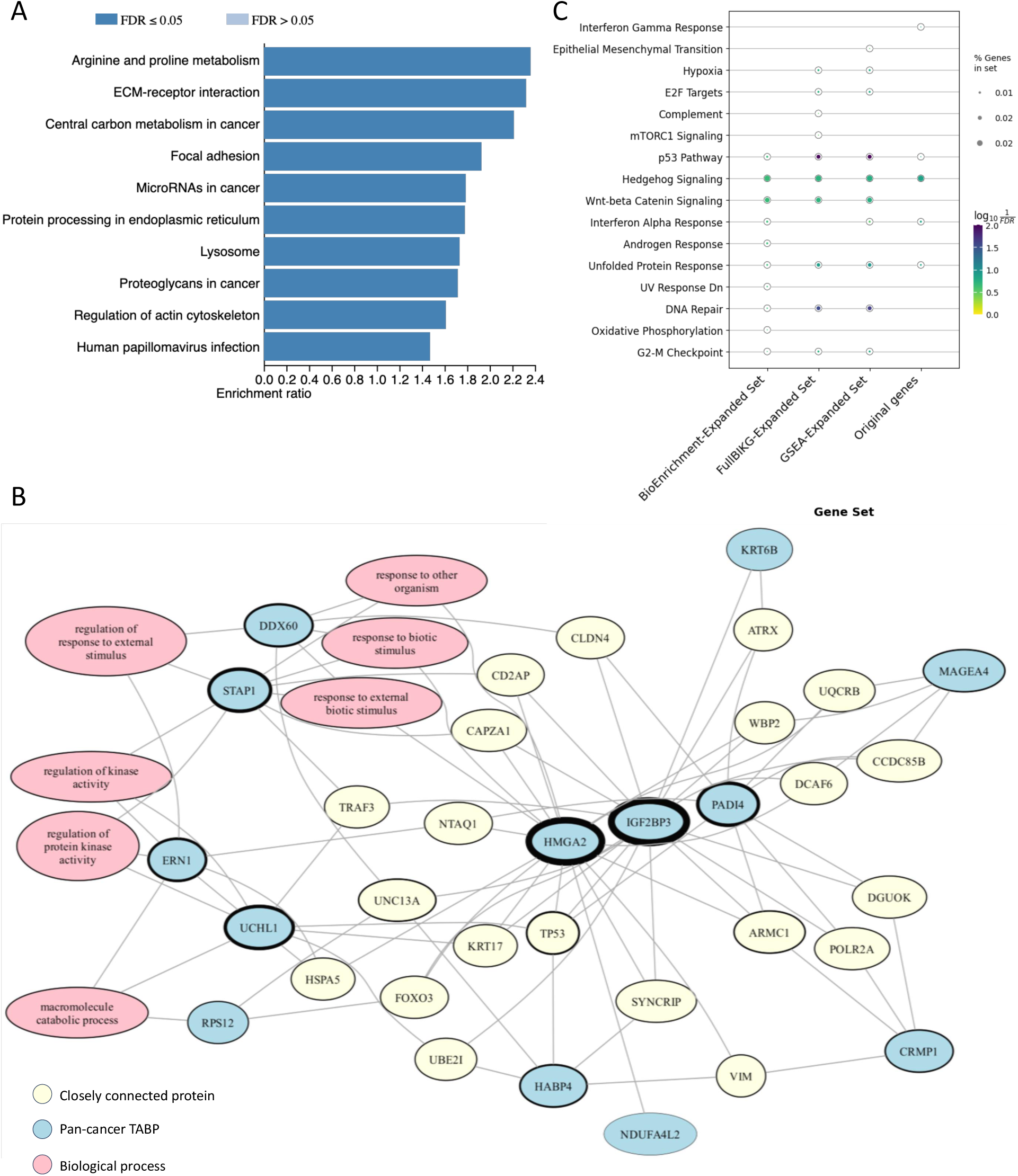
Signaling pathways associated with the TABPs. (A) Enrichment analysis of the tumor-associated bimodal proteins. (B) The subnetwork extracted from BIKG with 16 pan-cancer TABPs as seed nodes of PageRank. Blue nodes indicate the original TABPs; yellow, closely connected proteins to the TABPs; pink, biological processes. (C) Enrichment analysis using the BIKG-enriched proteins and original TABPs. The sizes of the colored circles represent the percentage of the gene covered in the gene set; colors represent –log10(FDR).

To explore the potential common characteristics of those TABPs, we investigated 16 pan-cancer TABPs (IGF2BP3, NDUFA4L2, EPS8L3, PADI4, UCHL1, TLNRD1, MSANTD2, CRMP1, ERN1, STAP1, RPS12, HABP4, KRT6B, MAGEA4, DDX60, HMGA2) that were commonly identified in at least three different indications (Supplementary Figure S4B, Supplementary Table S2). We reasoned that these 16 pan-cancer TABPs and their related pathways might reveal potential biomarkers. We utilized an AI-boosted knowledge graph, the Biological Insight Knowledge Graph (BIKG),^42^ and network propagation with the pan-cancer TABPs as seed nodes (Methods). In the BIKG-expanded network, we clustered 13 pan-cancer TABPs and 21 closely connected proteins (Figure 2B). Of the 21 proteins, 5 (KRT17, TP53, TRAF3, UBE2I, UNC13A) were identified as bimodal proteins in at least one tumor cohort in the CPTAC pan-cancer data sets (Supplementary Figure S4C–I), indicating that the AI-based knowledge graph could capture common signals on pan-cancer TABPs (chi-square test, *P* < 0.0001). Pathway enrichment analysis revealed a significant association of the proteins in the network with E2F targets, p53 pathway, WNT/β-catenin signaling, and DNA repair that were interconnected and played an important role in the development and progression of cancer^50–53^ (Figure 2C).

### TROP2 bimodality in COAD

TROP2, also known as tumor-associated calcium signal transducer 2 (TACSTD2), is a calcium signal transducer that is often overexpressed in many solid tumors, including colorectal cancer.^54,55^ Increased TROP2 expression is associated with tumor aggressiveness, epithelial-to-mesenchymal transition, metastasis, and decreased overall survival.^54,56,57^ These characteristics have triggered an interest in developing TROP2-targeted therapies for patients with colorectal cancer.^58^ In our study, TROP2 was identified as one of the top TABPs in COAD, leading us to further explore TROP2 bimodality to understand the mechanisms underlying this biological status.

TROP2 (TACSTD2) was reliably quantified in 41 tumor samples in the COAD cohort and had a bimodal distribution in tumor samples (*BI*, 3.24; 95% confidence interval [CI], 1.74-3.96; *BIC diff*, -24.08; 95% confidence interval [CI], -42.19 to -11.18; Figure 3A) but not adjacent normal colon tissues. A comparison with RNA-seq data for the TROP2 mRNA transcript revealed that the bimodal distribution of TROP2 in COAD tumors was also present at the transcriptomic level (Figure 3A). TACSTD2 mRNA levels were also significantly higher for the TROP2-high samples than for the TROP2-low samples (Mann-Whitney U-test, *P* ≤ 10e-4; Supplementary Figure S5A), in agreement with the strong correlation between TROP2 protein and mRNA levels (Spearman correlation, *P* = 1.5e-15; Supplementary Figure S5B). To further validate the discovery of TROP2 bimodality in colorectal cancer, we explored two independent proteomics data sets, from the T1NxM0 colorectal cancer (CRC) cohort in Chinese population and The Cancer Genome Atlas (TCGA) colorectal cancer cohort, and showed that in both datasets, TROP2 had bimodal distributions in tumor samples (T1NxM0 CRC cohort^59^: *BI* = 3.10, 95% confidence interval [CI] = 0.71–9.05, *BIC diff* = –3.00, 95% CI = –44.5 to 9.0; TCGA CRC cohort^43^: *BI* = 1.41, 95% CI = 0.52–4.10, *BIC diff* = –66.6, 95% CI = –108.2 to –34.8; Supplementary Figures S5C, S5D).

**Figure 3.**
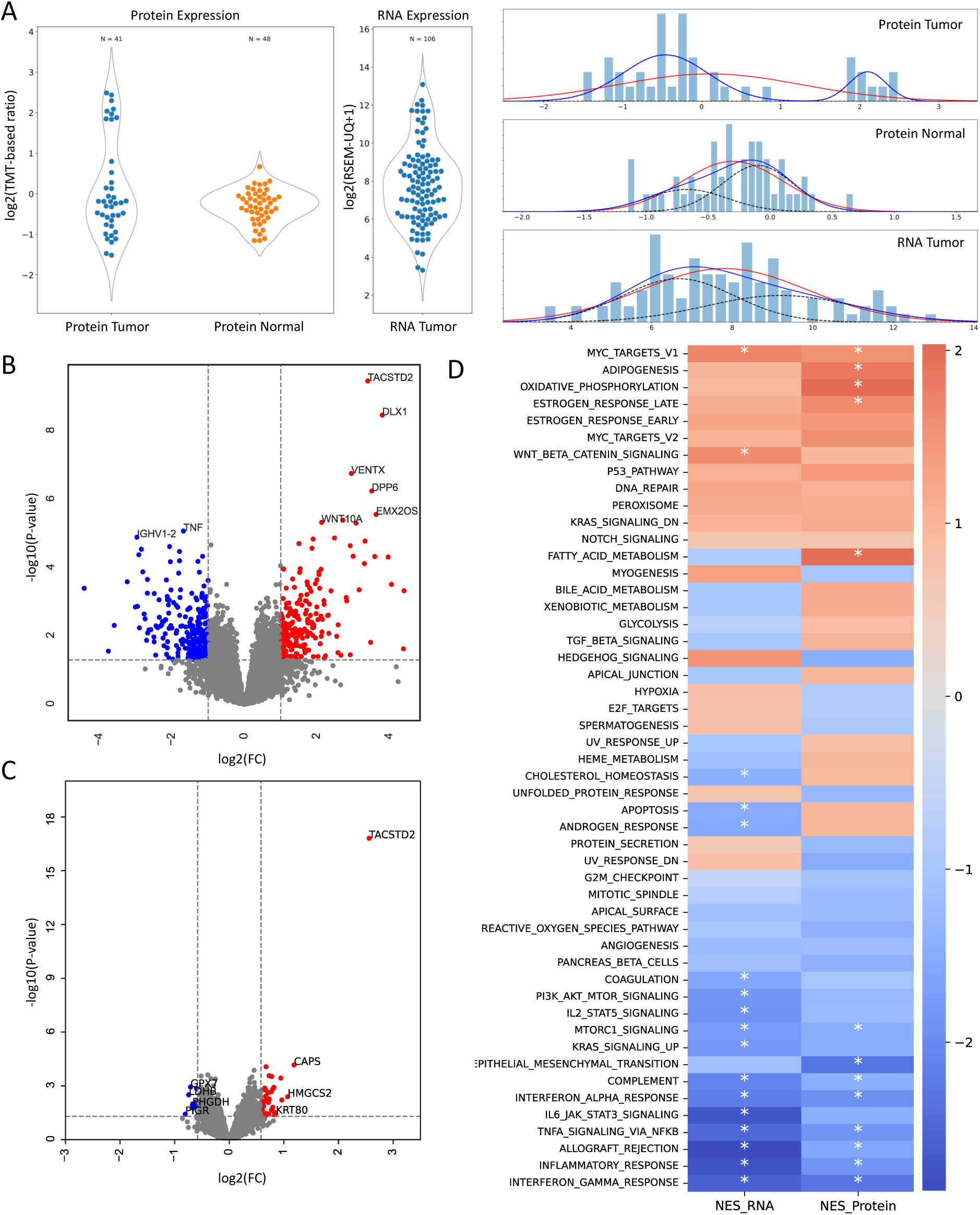
TROP2 (TACSTD2) bimodality in the CPTAC COAD. (A) Distributions of TROP2 protein expression in CPTAC COAD tumor and normal samples and RNA expression in tumors. Bimodality was identified only in proteomics data in tumor samples. The red line indicates the corresponding unimodal fit, and the blue and black lines show the bimodal fit. (B, C) Volcano plot showing differentially expressed genes/proteins between TROP2-high and TROP2-low samples in the CPTAC COAD cohort. Red indicates significantly upregulated genes (*P* < 0.05, log(FC) > 0.58); blue, significantly downregulated genes (*P* < 0.05, log(FC) < –0.58). (D) Heat map showing results of GSEA. Normalized enrichment scores are plotted for hallmark pathways from MSigDB based on differentially expressed genes and proteins (statistically significant results are indicated by an asterisk).

Next, we showed that TROP2 bimodality is not confounded by demographic and clinical features, chromosomal instability phenotype (CIN), microsatellite instability phenotype (MSI), or transcriptomic subtype (Supplementary Figures S6, S5E, S5F). Multi-omics factor analysis (MOFA)^60^ identified other features associated with TROP2 abundance. One of the latent factors fitted by the model had a high-weight contribution from both the TROP2 protein and mRNA abundance features but also from multiple copy number alteration features. Fisher exact test revealed that samples classified as TROP2-high were enriched for copy number amplifications of top genes associated with this latent factor (*P* = 0.006; Supplementary Figure S5G, Supplementary Table S2). *MYC* was identified as one of the top 10 genes associated with this latent factor through MOFA analysis and was a hub node in STRINGdb, connecting with 54 copy number alteration genes, including *VEGFR* and *CCNE1*/*CCNE2*. *VEGFR*, whose promoter is known to be targeted by *MYC*, is implicated in angiogenesis, while *CCNE1* and *CCNE2* play essential roles in G1/S cell cycle progression. However, statistical tests revealed that *MYC* copy number amplification was nonsignificant (*P* = 0.06; Supplementary Figure S5H), and *MYC* mRNA expression showed no significant difference between TROP2-high and TROP2-low samples (*P* = 0.74; Supplementary Figure S5I).

### Therapeutic opportunities driven by TROP2 bimodality

Differential RNA-seq/protein expression analysis between TROP2-high and TROP2-low samples using the DESeq2 package revealed a number of upregulated and downregulated genes/proteins (Figure 3B, 3C; Supplementary Table S2). Pathway enrichment for the differentially expressed genes and proteins revealed concordant enrichment of hallmark pathways, the most notable upregulated pathways in the TROP2-high group being the MYC pathway and WNT/β-catenin signaling pathways, through which TROP2 plays an important role in cell proliferation, apoptosis, and cell growth.^61–63^ In the TROP2-low group, inflammatory and interferon-related pathways, as well as tumor necrosis factor-alpha and IL6-JaAK-STAT3 signaling pathways, were upregulated (Figure 3D).

Phosphoproteomics analysis showed that in the TROP2-high samples, the most significantly upregulated phosphosites were EVPL_S2025, AXIN2_S70, NIPBL_T2667, and MYH14_S221 (Figure 4A). Kinase activity inference from phosphoproteomics identified some upregulated kinase activities in TROP2-high samples (false discovery rate, FDR > 0.05; Figure 4B). We utilized those kinases as seed nodes to identify a corresponding network in BIKG^42^ (Methods; Figure 4C), capturing associated proteins and biological processes. Pathway analysis based on this network revealed a correlation between the upregulated kinases and WNT/β-catenin signaling (Supplementary Figure S7A), consistent with prior differential analyses using proteomics/RNA-seq data.

**Figure 4.**
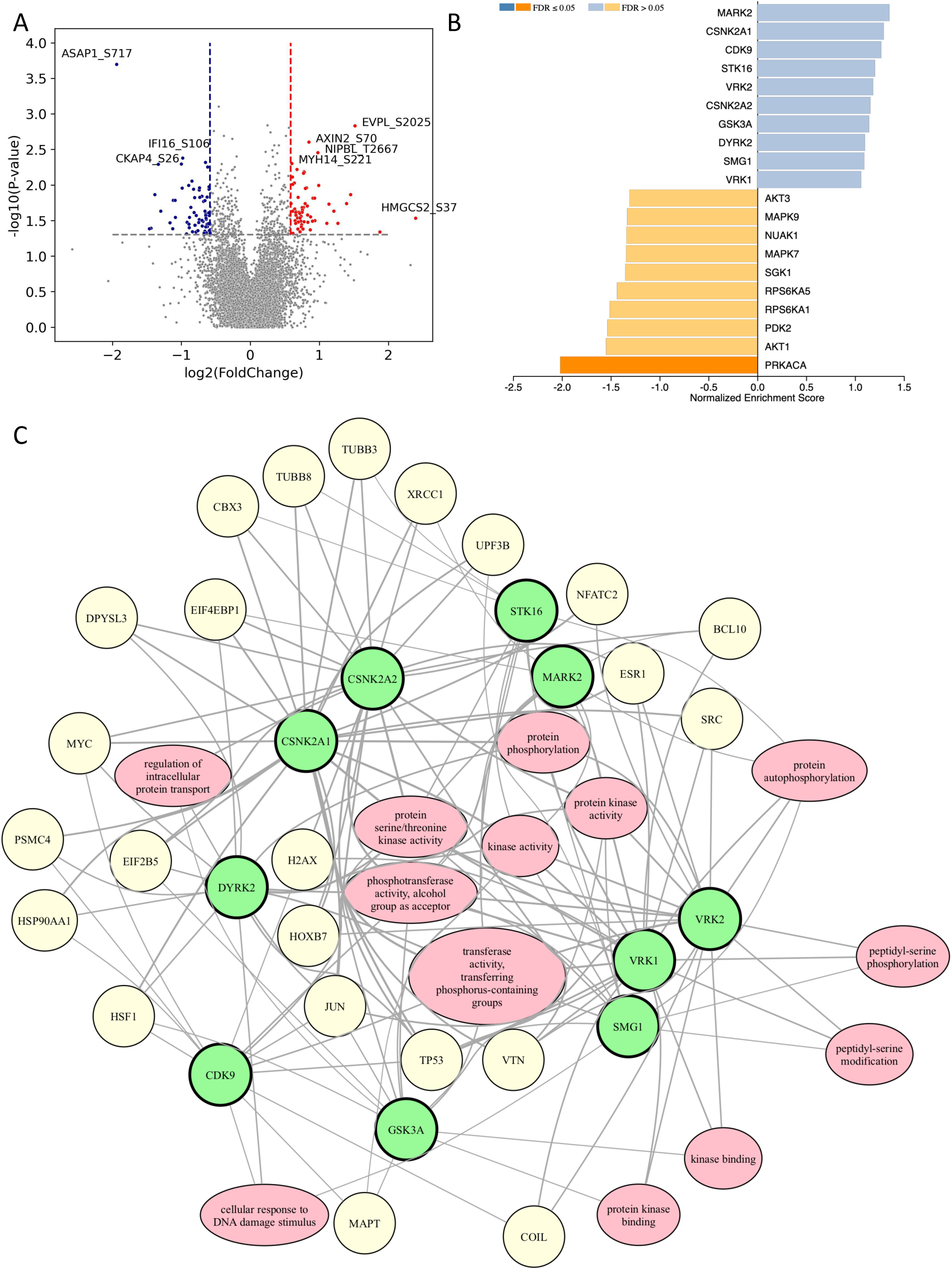
TROP2 bimodality and immune suppression in COAD. (A) Volcano plot showing differentially expressed phosphosites between TROP2-high and TROP2-low samples in the CPTAC COAD cohort. Red indicates significantly upregulated phosphosites (*P* < 0.05, LFC > 0.58); blue, significantly downregulated genes (*P* < 0.05, LFC < –0.58). (B) Kinase activity inferred from differentially expressed phosphosites using GSEA. (C) Network extracted from BIKG with upregulated kinases as seeds. Green nodes indicate the upregulated kinases; yellow, closely connected proteins to the upregulated kinases; pink, biological processes.

In contrast, kinase activity inference showed that TROP2-low samples had significantly higher PRKACA kinase activity (FDR < 0.05; Figure 4B). PRKACA protein expression was significantly correlated with the PI3K-AKT-MTOR signaling pathway in COAD (Supplementary Figure S7B). From BIKG-based network analysis, we identified upregulated immune-related pathways (Supplementary Figure S7C, S7D), which were also identified by the previous mRNA/protein DE analyses. Using the immune scores from ESTIMATE,^64^ we further confirmed that TROP2-low samples had more activated immune signaling than TROP2-high samples (Supplementary Figure S7E). The immune subtypes from a previous CPTAC study^65^ showed that TROP2-low samples are enriched with CD8-/interferon gamma-positive (IFNG^+^) subtypes as compared with CD8^−^/IFNG^−^ subtypes (chi-square test, *P* = 0.005; Supplementary Figure S7F). CD8^−^/IFNG^+^ represents an immune subtype with low immune infiltration of CD8 T cells and B cells, but strong activation of IFN signaling. CD8^−^/IFNG^−^ was low in both functions (immune cold).

## Discussion

We developed a bimodal model tailored for patient-derived proteomics data, which are often characterized by smaller sample sizes and more missing values than RNA-seq data. We used simulated bimodal and unimodal distributions across various sample sizes to provide reliable benchmark data for evaluating bimodality metrics. Theoretically, the *BI* and *BIC diff* metrics are very similar, in that they are derived from the same fitting procedure. A negative *BIC diff* indicates a bimodal preference, whereas a positive value suggests a unimodal fit. Unlike *BIC diff*, *BI* does not have defined thresholds and usually uses percentile thresholds or heuristic thresholds.^13,16,19,20^ Our results also indicated that smaller sample sizes affect the sensitivity and specificity of both metrics. We validated and established optimal thresholds for *BI* and *BIC diff* using a benchmarking data set that reflected actual sample conditions in proteomics data. This rigorous methodology ensured the robustness and reliability of our bimodality model, even under constrained sample conditions.

Our investigation into the bimodality of protein expressions within the CPTAC pan-cancer proteomics data has revealed insights into the molecular underpinnings of cancer biology. The identification of 2401 TABPs underscores the prevalence of bimodal expression in cancer and its potential utility in refining diagnostic and therapeutic strategies. Notably, our approach leverages advances in proteogenomics, addressing previous limitations inherent in RNA-based studies by directly incorporating protein expression data. This shift is crucial, given the often modest correlation between RNA and protein levels, which may not adequately reflect post-transcriptional modifications and the true biological state of the cells. For example, in contrast to findings in a previous pan-cancer RNA-seq–based bimodality study, we identified 2253 unique TABPs, of which 148 were overlapping.^20^

Our findings highlight the significant associations of TABPs with critical cancer pathways, such as arginine and proline metabolism, ECM-receptor interaction, and focal adhesion. These pathways are integral to tumor progression and metastasis, suggesting that proteins with bimodal expression patterns could serve as potential biomarkers for these biological processes in various tumor types. Moreover, the application of AI-boosted knowledge graphs in our analysis allowed for the identification of common patterns among these proteins across different cancers, potentially offering new avenues for targeted therapies.

TROP2 has been previously described as both a prognostic biomarker and a therapeutic target.^62^ Interestingly, our study identified two patient populations characterized by bimodal TROP2 expression. An in-depth multi-omics analysis revealed distinct disease biology associated with these populations, suggesting that TROP2 may serve as a novel biomarker for colon cancer. Specifically, MOFA analysis indicated that the TROP2-high group was strongly associated with the *MYC* signaling pathway, as revealed through differential RNA and protein pathway enrichment analysis. Despite this finding, statistical tests showed that *MYC* was not significantly copy number-amplified (*P* = 0.06; Supplementary Figure S5H), and its mRNA abundance did not differ between TROP2-high and TROP2-low samples (*P* = 0.74; Supplementary Figure S5I). This finding suggests that *MYC* pathway activation may occur through indirect mechanisms, potentially involving alternative regulatory elements or post-translational modifications.

In addition, RNA and protein analyses identified enrichment of the WNT/β-catenin pathway in the TROP2-high population. Previous studies have shown that TROP2 regulates self-renewal, proliferation, and transformation through regulated intramembrane proteolysis (RIP), which cleaves TROP2 into extracellular and intracellular domains.^62,66^ TROP2 can enhance the self-renewal of stem cells and progenitor cells via β-catenin signaling and can facilitate the development of precursor lesions in prostate cancer in vivo, with disruptions occurring in the absence of β-catenin or TROP2 RIP cleavage mutants.^62,66–70^ Our findings suggest that TROP2 may have an analogous role in colon cancers. The Colorectal Cancer Subtyping Consortium (CRCSC), a group that analyzed gene expression data in more than 30 expression sets spanning multiple platforms and sample preparation methods, has proposed consensus molecular subtypes (CMSs) based on genome-wide gene expression profiling analysis, which categorizes colorectal cancers into four groups (CMS1–4).^71,72^ TROP2-high colorectal cancer tumors are enriched in the CMS2 subtype of colorectal cancer, which is also characterized by WNT/β-catenin activation.^72^ Furthermore, phosphoproteomics analysis revealed a significant increase in AXIN2 phosphorylation. As an inhibitor of the WNT/β-catenin pathway, AXIN2 facilitates the phosphorylation of β-catenin and adenomatous polyposis of the colon by GSK3B^73,74^ and promotes the degradation of β-catenin.^75^ This phenomenon may represent a negative feedback loop through AXIN2 to control β-catenin activation in colorectal cancer, potentially driven by TROP2 expression.^73^

In contrast, TROP2-low populations are distinguished by activated PI3K-AKT pathways and inflammatory and interferon-related pathways, as demonstrated by differential RNA and protein expression pathway enrichment analysis. Phosphoproteomics analysis showed that TROP2-low samples had significantly higher PRKACA kinase activity (FDR < 0.05; Figure 4B). PRKACA protein expression significantly correlated with the PI3K-AKT-MTOR signaling pathway in COAD (Supplementary Figure S7B), suggesting that in TROP2-low samples, PRKACA could drive colorectal cancer progression via the PKA-FAK-AKT pathway.

These findings suggest that oncogenesis and tumor proliferation in the two populations are potentially driven by distinct signaling pathways and possess different tumor microenvironments. Therefore, it is crucial to consider therapeutic interventions guided by TROP2 protein expression as a biomarker. This personalized approach could enhance treatment efficacy by targeting the specific molecular drivers present in each tumor subtype. For instance, if the role of TROP2 in self-renewal and transformation is validated in the TROP2-high colorectal cancer population, therapeutics that block TROP2 activation might constitute an effective treatment strategy for this group. Further experiments are warranted to test this compelling hypothesis.

Despite these advancements, our study has limitations, including its reliance on available proteomics data that may not include all cancer types or all stages of disease progression. CPTAC has insufficient therapeutic information to uncover biomarkers with our methods. In addition, although we demonstrated the application of our bimodality model to a large data set, the generalizability of these findings to other data sets or through prospective validation remains to be tested. One example we showed is TROP2 bimodality validation in two external data sets.

Future research should focus on expanding the application of bimodality analyses to other types of cancer and integrating multi-omics data to further validate and refine the identified biomarkers. Moreover, clinical trials could explore the therapeutic implications of these findings, particularly the differential strategies suggested by TROP2 bimodality in colorectal cancer.

In conclusion, our study not only provides a robust framework for the identification of bimodal proteins expressed across cancers but also highlights the role of such biomarkers in elucidating the complex molecular landscape of malignancies, offering promising prospects for precision oncology.

## Methods

### Data acquisition

CPTAC processed RNA-seq, proteomics, phosphoproteomics and clinical data were downloaded from LinkedOmics.^77^ The clinical data used in the portal were collected from CPTAC with the May 2022 update. Age was truncated to 90 years. Tumors with a size of ≤0 cm were replaced with “NA” (not applicable). CPTAC immune-related information was downloaded from Petralia et al.^65^

TCGA processed proteomics data from colon and rectal cancer were downloaded as described Zhang et al.^43^ Processed proteomics data from T1NxM0 colorectal cancer were downloaded as described in Zhuang et al.^59^

### Simulated bimodality distributions

Data simulation was carried out by creating linear combinations of two normal distributions. The number of bimodal genes was controlled by sampling a binomial distribution. For the genes chosen as bimodal, a weight was randomly sampled. The difference in means between the modes of the normals was randomly sampled from a uniform distribution, assigning the value to the first mode and 0 to the second. If the genes were selected as unimodal, then both means were assigned as equal. Finally, the variance was kept equal for both modes and randomly chosen from a uniform distribution. See “Benchmarking *BI* and *BIC diff*” below for mode details.

### Bimodality metrics

The approach used to detect proteins with bimodal expression entailed determining whether the distribution is best described as unimodal or bimodal. Technically, the approach involved either (1) fitting the data to these two models and using goodness-of-fit to make a choice or (2) using a metric that transitions to different values if the data are unimodal or bimodal. Both the *BI* and the *BIC diff* belong to the first class, whereas measuring *kurt* belongs to the second one. For each simulation data, the *kurt* of the data was computed directly from the samples. The *BI* and the *BIC diff* were computed after fitting a normal (unimodal) or a mixture of two normal (bimodal) distributions, indicated by the respective red and blue lines in Supplementary Figure S1.

To detect proteins or genes with bimodal expression, three different metrics were computed (Supplementary Table S1). The first metric was used to obtain the *kurt* of the distribution that measures the “tailness,” by computing:

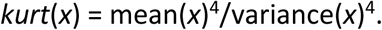

The second metric was used to determine the difference in goodness-of-fit (measured by the *BIC*) when the data were fitted using a normal distribution or a mixture of normal distributions, by computing:

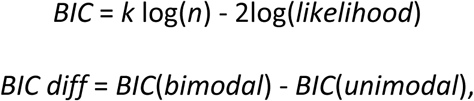

where *likelihood* is the maximized value of the likelihood for the model, *n* is the number of data points, *k* is the number of parameters, *BIC*(*bimodal*) is the score obtained upon fitting a weighted mixture of normal distributions, and *BIC*(*unimodal*) is the score obtained by fitting a normal distribution. Negative values suggest that the data are better modeled as a mixture of normals, whereas positive values indicate a unimodal distribution.

The third metric reused the models fitted above and computed the *BI*:

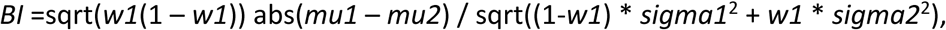

where *w1*, *mu1*, *sigma1*, *mu2*, and *var2* are the bimodal distribution fitted:

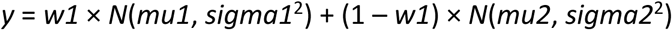

and *N*(*mu1*, *sigma1*^2^) is Gaussian. In general, *BI* is higher for distribution showing bimodality. This can be observed directly in the formula above. If the weights for the two modes are similar, *w1*\*(1 – *w1*) is higher. If the separation for the modes is high, then abs(*mu1* – *mu2*) is higher. If the spread of each mode is lower, then the denominator is smaller, and *BI* is larger.

One potential challenge posed by this type of analysis is how to assess the confidence of the metrics when the number of points used to fit the distribution is small. To overcome this issue, the samples for each protein analyzed were bootstrapped 300 times and the corresponding 95% CIs were computed.

### Benchmarking *BI* and *BIC diff*

*BI* and *BIC diff* metrics are very similar, given that both arise from the same fitting procedure. To assess the performance of these metrics, we created a set of known bimodal and unimodal distributions (*N* = 100). The number of bimodal and unimodal distributions was randomly chosen from a binomial distribution with *P* = 0.5. To completely define each distribution, we randomly sampled the following parameters:

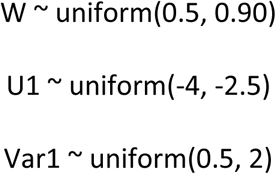

If the distribution was unimodal, then we created data following N(U1, Var1); if the distribution was bimodal, we created data following w*N(u1, var1) + (1-w1)*N(0, var1).

### Bimodality identification using CPTAC pan-cancer proteomics data sets

Ten tumor cohorts and six normal cohorts in CPTAC pan-cancer data sets (with at least 40 samples in the proteomics data) were analyzed. For each cohort, we calculated *BI* and *BIC diff* for each protein that was quantified in at least 30 samples in the cohort (Supplementary Table S1). To define bimodal proteins, we selected proteins according to the thresholds in Table 1.

TABPs were defined as those proteins identified as bimodal proteins in a tumor cohort but not in the matched normal tissues.

### Prioritizing proteins of interest using BIKG

Biological understanding of TABPs or differential proteins was identified using knowledge graph network analysis.^42^ Knowledge graph data structures provide flexibility in network modeling by providing variants of both edge and node types. In this study, we used this analysis for two cases: (1) expanding gene sets of interest to identify related gene targets and (2) using biologically relevant node types (genes, pathways, molecular functions, disease, and biological processes) to find underlying biology to describe the relationships between our target genes.

The two knowledge graphs were constructed using data from the publicly available databases METABASE, DOROTHEA, HETIONET, and CELLPHONEDB. In addition, relations between gene nodes in both subgraphs were filtered to contain edge relations of types: interacts, regulates, or modifies. Relations between gene targets and other node types in the biological enrichment graph were of the type “participates” for biological process, pathway, and molecular function node types, and “up/downregulates” for connections with disease nodes. For each gene set of interest, the two subgraphs were further filtered to retain edges connecting either genes with this list itself or connecting any node in this set to any other node in the graph. Such filtering provided a more manageable size of network to analyze.

A repeated random walk procedure was performed where *N* = 1000 random paths were traversed originating from each node in our target gene list. Upon performing all the walks, edges that had a number of occurrences above a certain threshold were retained. This random walk procedure was repeated until the obtained network did not change between iterations, providing network stability. Such subgraphs generated for each gene list of interest provided expanded gene signatures of interest as well as biological events that may relate to our genes. The expanded gene sets were then used in gene set enrichment analysis (GSEA) to identify additional pathways related to the larger gene set expanded using the knowledge graph approach.

### Transcriptomic subtypes in COAD

The transcriptomic subtype classification of each sample was performed according to the Consensus Molecular Subtypes framework as described in Guinney et al.^71^

### Microsatellite instability status and CIN in COAD

Both MSI status and CIN information was obtained from LinkedOmics according to detailed methods described by Vasaikar et al.^77^

### Statistical hypothesis test

The chi-square test of independence was used to assess whether there was a significant association between the CMS transcriptomic subtypes/demographic/clinical features/MSI/CIN and TROP2 high versus low status.

### Multifactor omics integration using MOFA+

MOFA^78^ is a latent variable model that relies on matrix factorization and Bayesian modeling to uncover orthogonal axes of heterogeneity in multimodal data sets. The dependencies between omics layers are modeled by decomposing data into a lower-dimensional representation of shared factors and data set–specific contributions. This approach enables the identification of molecular signatures that are concordant across different omics modalities while accounting for data set–specific variation. In our study, MOFA was employed to determine which omics features, including proteomics, transcriptomics, copy number variations, miRNAs, mutations, and phospho-proteomics, were associated with the bimodal TROP2 protein distribution in the CPTAC COAD patient cohort. Features with the highest weight in the MOFA factor that also had a high-weight contribution from TROP2 protein were selected for downstream investigation.

### Differential analysis (RNA-seq, proteomics, and phosphoproteomics)

Tumor samples from the COAD cohort for RNA-seq, proteomics, and phosphoproteomics were used for differential expression analysis between TROP2-high and TROP2-low subgroups.

For RNA-seq, differential expression analysis was performed using the Python implementation of the DESeq2^79,80^ (version 0.3.1) package in R (version 1.34.0). Briefly, raw count data for 60,660 gene transcripts were loaded into Python, and low-count genes were filtered out to minimize noise (requiring that on average, each transcript has been detected 10 times in each sample), yielding a total analysis set of 19,419 reliably expressed transcripts. Normalization factors were calculated to account for differences in library size, and the negative binomial distribution was fitted to the count data. Dispersion estimation and Wald tests were performed to identify genes that were differentially expressed between the TROP2-high and TROP2-low samples. To correct for multiple hypothesis testing, *P* values were adjusted using the Benjamini-Hochberg method.^81^ Genes with an adjusted *P* < 0.05 were considered to be significantly differentially expressed.

For proteomics, proteins that were quantified in at least 50% of the samples were used for further analysis. Imputation and differential protein expression analysis was performed using DEP package in the R version 4.3.1 environment as described in Wang et al.^41^ The fgsea package^82^ was used for protein gene set enrichment analysis. The fgsea algorithm was run with 1000 permutations.

For phosphoproteomics, a phosphosite must have been quantified in at least 50% of tumor samples or was removed. The unpaired Wilcoxon rank-sum test was used to calculate significance. Significantly differential sites were defined as *P* < 0.05 and Fold Change, *FC* > 1.5.

### GSEA

The GSEA “prerank” method^83^ was employed to assess pathway enrichment, using a list of genes ranked by *P* values from differential RNA/protein analysis or copy number variations associated with MOFA latent factors. GSEA was used to evaluate whether predefined gene sets (“hallmarks” or “C1 positional gene sets corresponding to human chromosome cytogenic bands”) from the MSigDB database were significantly enriched toward the top (positively enriched) or the bottom (negatively enriched) of the ranked gene/protein list. Statistical significance was determined through a permutation-based approach that computed enrichment scores and false discovery rates.

### Kinase-substrate association database

We constructed the gmt files for GSEA to infer kinase activity. The ground truth kinase substrate associations were downloaded from GPS 5.0,^83^ a curation of 15,194 experimentally identified kinase substrate associations. The prediction kinase substrate associations were obtained from CoPheeKSA.^84^ When a kinase had more than 100 known substrates, no predicted substrates were added to this kinase. When a kinase had fewer than 100 known substrates, the predicted substrates from CoPheeKSA were added to this kinase, with a maximum number of 500 (ranked by predictions scores).

### Kinase activity inference

We used Webgestalt^85^ to perform GSEA on the phosphosites and the –log10(*P*) from differential analysis.^86^ The organism was set to “other.” The functional database was uploaded using a self-defined gmt file (kinase-substrate association [KSA] database). The output is the kinase activity inferred from phosphoproteomics data.

## Data availability

For CPTAC BRCA, ccRCC, COAD, GBM, HNSCC, LSCC, LUAD, OV, PDAC and UCEC, processed RNA-Seq, proteomics, phosphoproteomics and clinical data matrices were downloaded from LinkedOmics^77^: https://www.linkedomics.org/login.php.

## Acknowledgments

We gratefully acknowledge the CPTAC for providing open-source proteomics data and Deborah Shuman of AstraZeneca for editing the article and formatting the figures. This study was funded by AstraZeneca.

## Author contributions

Conceptualization, J.W. and W.Z.; Methodology, W.J., D.B. and W.Z.; Formal Analysis, W.J., D.B., J.Z., J.W., S.J., Y.L.; Investigation, W.J., M.S., J.H., J.Z., W.Z.; Data Curation, J.W; Writing – Original Draft, W.J., D.B, J.Z., J.W.; Visualization, W.J., D.B., J.Z., J.W., S.J.; Review and editing, M.S., J.H., J.Z., K.B., W.Z.

## References

1 Cancer Genome Atlas Network. Comprehensive molecular portraits of human breast tumours. Nature 490, 61–70 (2012).

2 Garnett, M. J. et al. Systematic identification of genomic markers of drug sensitivity in cancer cells. Nature 483, 570–575 (2012).

3 Mertins, P. et al. Proteogenomics connects somatic mutations to signalling in breast cancer. Nature 534, 55–62 (2016).

4 Sarhadi, V. K. & Armengol, G. Molecular biomarkers in cancer. Biomolecules 12, 1021 (2022).

5 Schwartzberg, L., Kim, E. S., Liu, D. & Schrag, D. Precision oncology: who, how, what, when, and when not? Am Soc Clin Oncol Educ Book 37, 160–169 (2017).

6 Yang, W. et al. Genomics of drug sensitivity in cancer (GDSC): a resource for therapeutic biomarker discovery in cancer cells. Nucleic Acids Res 41, D955–D961 (2013).

7 Davies, H. et al. Mutations of the BRAF gene in human cancer. Nature 417, 9499–9454 (2002).

8 Soda, M. et al. Identification of the transforming EML4-ALK fusion gene in non-small-cell lung cancer. Nature 448, 561–566 (2007).

9 Tomlins, S. A. et al. Recurrent fusion of TMPRSS2 and ETS transcription factor genes in prostate cancer. Science 310, 644–648 (2005).

10 Vasaikar, S. et al. Proteogenomic analysis of human colon cancer reveals new therapeutic opportunities. Cell 177, 1035–1049 (2019).

11 Dagogo-Jack, I. & Shaw, A. T. Tumour heterogeneity and resistance to cancer therapies. Nat Rev Clin Oncol 15, 81–94 (2018).

12 Justino, J. R., Reis, C. F. D., Fonseca, A. L., Souza, S. J. & Stransky, B. An integrated approach to identify bimodal genes associated with prognosis in cancer. Genet Mol Biol 44, e20210109 (2021).

13 Moody, L., Mantha, S., Chen, H. & Pan, Y. X. Computational methods to identify bimodal gene expression and facilitate personalized treatment in cancer patients. J Biomed Inform 100S, 100001 (2019).

14 Kim, C. et al. Estrogen receptor (ESR1) mRNA expression and benefit from tamoxifen in the treatment and prevention of estrogen receptor-positive breast cancer. J Clin Oncol 29, 4160–4167 (2011).

15 Muftah, A. A. et al. Further evidence to support bimodality of oestrogen receptor expression in breast cancer. Histopathology 70, 456–465 (2017).

16 Ertel, A. Bimodal gene expression and biomarker discovery. Cancer Inform 9, 11–14 (2010).

17 Tong, P., Chen, Y., Su, X. & Coombes, K. R. SIBER: systematic identification of bimodally expressed genes using RNAseq data. Bioinformatics 29, 605–613 (2013).

18 de Torrenté, L. et al. The shape of gene expression distributions matter: how incorporating distribution shape improves the interpretation of cancer transcriptomic data. BMC Bioinformatics 21, 562 (2020).

19 Ochab-Marcinek, A. & Tabaka, M. Bimodal gene expression in noncooperative regulatory systems. Proc Natl Acad Sci U S A 107, 22096–22101 (2010).

20 Ba-Alawi, W. et al. Bimodal gene expression in patients with cancer provides interpretable biomarkers for drug sensitivity. Cancer Res 82, 2378–2387 (2022).

21 Gong, Y. et al. Determination of oestrogen-receptor status and ERBB2 status of breast carcinoma: a gene-expression profiling study. Lancet Oncol 8, 203–211 (2007).

22 Kernagis, D. N., Hall, A. H. & Datto, M. B. Genes with bimodal expression are robust diagnostic targets that define distinct subtypes of epithelial ovarian cancer with different overall survival. J Mol Diagn 14, 214–222 (2012).

23 Rimm, D. L. et al. Bimodal population or pathologist artifact? J Clin Oncol 25, 2487–2488 (2007).

24 Schnitt, S. J. Estrogen receptor testing of breast cancer in current clinical practice: what’s the question? J Clin Oncol 24, 1797–1799 (2006).

25 Bessarabova, M. et al. Bimodal gene expression patterns in breast cancer. BMC Genomics 11 Suppl 1, S8 (2010).

26 Mason, C. C. et al. Bimodal distribution of RNA expression levels in human skeletal muscle tissue. BMC Genomics 12, 98 (2011).

27 Liu, Y., Beyer, A. & Aebersold, R. On the dependency of cellular protein levels on mRNA abundance. Cell 165, 535–550 (2016).

28 Nie, L., Wu, G. & Zhang, W. Correlation of mRNA expression and protein abundance affected by multiple sequence features related to translational efficiency in Desulfovibrio vulgaris: a quantitative analysis. Genetics 174, 2229–2243 (2006).

29 Cao, L. et al. Proteogenomic characterization of pancreatic ductal adenocarcinoma. Cell 184, 5031–5052. (2021).

30 Clark, D. J. et al. Integrated proteogenomic characterization of clear cell renal cell carcinoma. Cell 179, 964–983 (2019).

31 Dong, L. et al. Proteogenomic characterization identifies clinically relevant subgroups of intrahepatic cholangiocarcinoma. Cancer Cell 40, 70–87 (2022).

32 Geffen, Y. et al. Pan-cancer analysis of post-translational modifications reveals shared patterns of protein regulation. Cell 186, 3945–3967 (2023).

33 Krug, K. et al. Proteogenomic landscape of breast cancer tumorigenesis and targeted therapy. Cell 183, 1436–1456 (2020).

34 Li, Y. et al. Proteogenomic data and resources for pan-cancer analysis. Cancer Cell 41, 1397–1406 (2023).

35 Liao, Y. et al. A proteogenomics data-driven knowledge base of human cancer. Cell Syst 14, 777–778 (2023).

36 Mani, D. R. et al. Cancer proteogenomics: current impact and future prospects. Nat Rev Cancer 22, 298–313 (2022).

37 Ng, C. K. Y. et al. Integrative proteogenomic characterization of hepatocellular carcinoma across etiologies and stages. Nat Commun 13, 2436 (2022).

38 Wang, L. B. et al. Proteogenomic and metabolomic characterization of human glioblastoma. Cancer Cell 39, 509–528 (2021).

39 Zhang, H. et al. Integrated proteogenomic characterization of human high-grade serous ovarian cancer. Cell 166, 755–765 (2016).

40 Zhang, Y., Chen, F., Chandrashekar, D. S., Varambally, S. & Creighton, C. J. Proteogenomic characterization of 2002 human cancers reveals pan-cancer molecular subtypes and associated pathways. Nat Commun 13, 2669 (2022).

41 Wang, J. et al. Pan-cancer proteomics analysis to identify tumor-enriched and highly expressed cell surface antigens as potential targets for cancer therapeutics. Mol Cell Proteomics 22, 100626 (2023).

42 Gogleva, A. et al. Knowledge graph-based recommendation framework identifies drivers of resistance in EGFR mutant non-small cell lung cancer. Nat Commun 13, 1667 (2022).

43 Zhang, B. et al. Proteogenomic characterization of human colon and rectal cancer. Nature 513, 382–387 (2014).

44 Huang, C. et al. Proteogenomic insights into the biology and treatment of HPV-negative head and neck squamous cell carcinoma. Cancer Cell 39, 361–379 (2021).

45 Satpathy, S. et al. A proteogenomic portrait of lung squamous cell carcinoma. Cell 184, 4348–4371 (2021).

46 Gillette, M. A. et al. Proteogenomic characterization reveals therapeutic vulnerabilities in lung adenocarcinoma. Cell 182, 200–225 (2020).

47 Dou, Y. et al. Proteogenomic characterization of endometrial carcinoma. Cell 180, 729–748 (2020).

48 Majo, S., Courtois, S., Souleyreau, W., Bikfalvi, A. & Auguste, P. Impact of extracellular matrix components to renal cell carcinoma behavior. Front Oncol 10, 625 (2020).

49 Geiger, B. & Yamada, K. M. Molecular architecture and function of matrix adhesions. Cold Spring Harb Perspect Biol 3, a005033 (2011).

50 Chen, H. Z., Tsai, S. Y. & Leone, G. Emerging roles of E2Fs in cancer: an exit from cell cycle control. Nat Rev Cancer 9, 785–797 (2009).

51 Groelly, F. J., Fawkes, M., Dagg, R. A., Blackford, A. N. & Tarsounas, M. Targeting DNA damage response pathways in cancer. Nat Rev Cancer 23, 78–94 (2023).

52 Wang, H., Guo, M., Wei, H. & Chen, Y. Targeting p53 pathways: mechanisms, structures, and advances in therapy. Signal Transduct Target Ther 8, 92 (2023).

53 Yu, F. et al. Wnt/beta-catenin signaling in cancers and targeted therapies. Signal Transduct Target Ther 6, 307 (2021).

54 Fang, Y. J. et al. Elevated expressions of MMP7, TROP2, and survivin are associated with survival, disease recurrence, and liver metastasis of colon cancer. Int J Colorectal Dis 24, 875–884 (2009).

55 Švec, J. et al. TROP2 represents a negative prognostic factor in colorectal adenocarcinoma and its expression is associated with features of epithelial-mesenchymal transition and invasiveness. Cancers (Basel) 14, 4137 (2022).

56 Fong, D. et al. High expression of TROP2 correlates with poor prognosis in pancreatic cancer. Br J Cancer 99, 1290–1295 (2008).

57 Hsu, E. C. et al. Trop2 is a driver of metastatic prostate cancer with neuroendocrine phenotype via PARP1. Proc Natl Acad Sci U S A 117, 2032–2042 (2020).

58 Alese, O. B. et al. Update on emerging therapies for advanced colorectal cancer. Am Soc Clin Oncol Educ Book 43, e389574 (2023).

59 Zhuang, A. et al. Proteomic characteristics reveal the signatures and the risks of T1 colorectal cancer metastasis to lymph nodes. Elife 12, e82959 (2023).

60 Argelaguet, R. et al. Multi-omics factor analysis: a framework for unsupervised integration of multi-omics data sets. Mol Syst Biol 14, e8124 (2018).

61 Lin, J. C. et al. TROP2 is epigenetically inactivated and modulates IGF-1R signalling in lung adenocarcinoma. EMBO Mol Med 4, 472–485 (2012).

62 Shvartsur, A. & Bonavida, B. Trop2 and its overexpression in cancers: regulation and clinical/therapeutic implications. Genes Cancer 6, 84–105 (2015).

63 Stepan, L. P. et al. Expression of Trop2 cell surface glycoprotein in normal and tumor tissues: potential implications as a cancer therapeutic target. J Histochem Cytochem 59, 701–710 (2011).

64 Yoshihara, K. et al. Inferring tumour purity and stromal and immune cell admixture from expression data. Nat Commun 4, 2612 (2013).

65 Petralia, F. et al. Pan-cancer proteogenomics characterization of tumor immunity. Cell 187, 1255–1277 (2024).

66 Stoyanova, T. et al. Regulated proteolysis of Trop2 drives epithelial hyperplasia and stem cell self-renewal via beta-catenin signaling. Genes Dev 26, 2271–2285 (2012).

67 Bignotti, E. et al. Trop-2 protein overexpression is an independent marker for predicting disease recurrence in endometrioid endometrial carcinoma. BMC Clin Pathol 12, 22 (2012).

68 Cubas, R., Li, M., Chen, C. & Yao, Q. Trop2: a possible therapeutic target for late stage epithelial carcinomas. Biochim Biophys Acta 1796, 309–314 (2009).

69 Cubas, R., Zhang, S., Li, M., Chen, C. & Yao, Q. Trop2 expression contributes to tumor pathogenesis by activating the ERK MAPK pathway. Mol Cancer 9, 253 (2010).

70 Liu, X. et al. Advances in Trop2-targeted therapy: novel agents and opportunities beyond breast cancer. Pharmacol Ther 239, 108296 (2022).

71 Guinney, J. et al. The consensus molecular subtypes of colorectal cancer. Nat Med 21, 1350–1356 (2015).

72 Li, X. et al. A modified protein marker panel to identify four consensus molecular subtypes in colorectal cancer using immunohistochemistry. Pathol Res Pract 220, 153379 (2021).

73 Jho, E. H. et al. Wnt/beta-catenin/Tcf signaling induces the transcription of Axin2, a negative regulator of the signaling pathway. Mol Cell Biol 22, 1172–1183 (2002).

74 Wu, Z. Q. et al. Canonical Wnt suppressor, Axin2, promotes colon carcinoma oncogenic activity. Proc Natl Acad Sci U S A 109, 11312–11317 (2012).

75 Bernkopf, D. B., Bruckner, M., Hadjihannas, M. V. & Behrens, J. An aggregon in conductin/axin2 regulates Wnt/beta-catenin signaling and holds potential for cancer therapy. Nat Commun 10, 4251 (2019).

76 National Cancer Institute. Proteomic Data Commons. https://pdc.cancer.gov/pdc/cptac-pancancer. Accessed February 13, 2025.

77 Vasaikar, S. V., Straub, P., Wang, J. & Zhang, B. LinkedOmics: analyzing multi-omics data within and across 32 cancer types. Nucleic Acids Res 46, D956–D963 (2018).

78 Argelaguet, R. et al. MOFA+: a statistical framework for comprehensive integration of multi-modal single-cell data. Genome Biol 21, 111 (2020).

79 Love, M. I., Huber, W. & Anders, S. Moderated estimation of fold change and dispersion for RNA-seq data with DESeq2. Genome Biol 15, 550 (2014).

80 Muzellec, B., Telenczuk, M., Cabeli, V. & Andreux, M. PyDESeq2: a python package for bulk RNA-seq differential expression analysis. Bioinformatics 39, btad547 (2023).

81 Benjamini, Y. & Hochberg, Y. Controlling the false discovery rate: a practical and powerful approach to multiple testing. J R Stat Soc Ser B 57, 289–300 (1995).

82 Korotkevich, G., et al. Fast gene set enrichment analysis. bioRxiv (2021). https://www.biorxiv.org/content/10.1101/060012v3. Accessed February 11, 2025.

83 The Cuckoo Workgroup. Group-based prediction system. http://gps.biocuckoo.cn/ (2025). Accessed February 13, 2025.

84 Jiang, W., et al. Illuminating the dark cancer phosphoproteome through a machine-learned co-regulation map of 26,280 phosphosites. (2024). https://www.biorxiv.org/content/10.1101/2024.03.19.585786v1. Accessed February 11, 2025.

85 Elizarraras, J., Liao, Y., Shi, Z. & Zhang, B. WEB-based Gene SeT AnaLysis Toolkit. https://www.webgestalt.org/. Accessed February 13, 2025.

86 Liao, Y., Wang, J., Jaehnig, E. J., Shi, Z. & Zhang, B. WebGestalt 2019: gene set analysis toolkit with revamped UIs and APIs. Nucleic Acids Res 47, W199–W205 (2019).

