## Supplementary Figure legends for "Bimodality in pan-cancer proteomics reveals new opportunities for biomarker discovery"

Bulusu K, Zhong W

**Supplementary Figure**

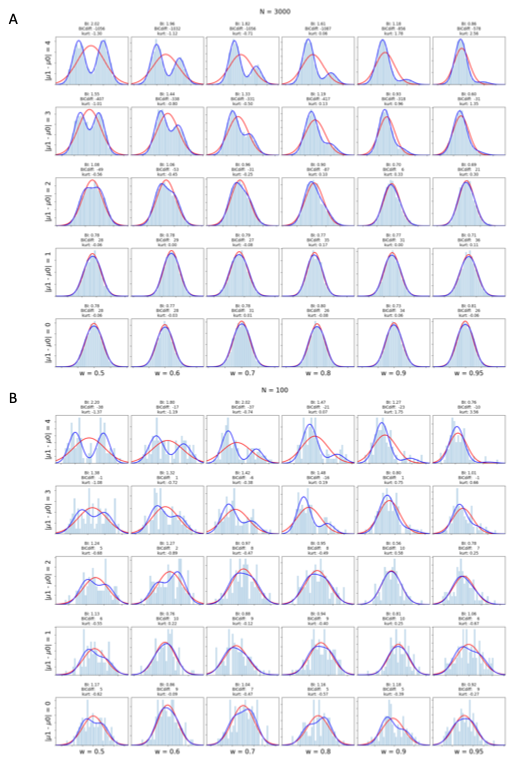

**Supplementary Figure S1.** Effect of changing the relative weights and mean separation for a mixture of Gaussian used for the bimodality model for proteomics. The y-axis shows what happens upon changing the difference in means of the two distributions (the variance is equal and kept at 1 for all cases). The x-axis shows the effect of changing the weight of the linear combination. (A) Three thousand samples were drawn randomly from each model; the red and blue lines indicate the corresponding unimodal and bimodal fits, respectively. (B) Same combinations as in (A) but on only 100 samples. Three bimodality metrics (bimodality index, *BI*; the difference in the Bayesian information criteria, *BIC diff*; and kurtosis, *kurt*) were calculated for each sample case.

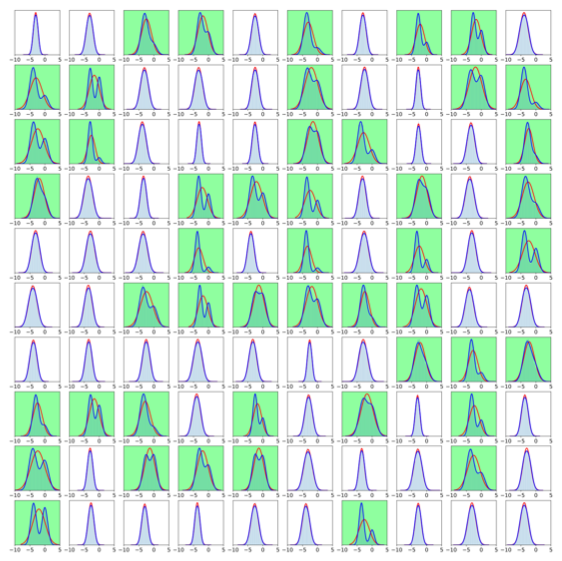
**Supplementary Figure S2.** Benchmarking data set of 100 known bimodal and unimodal distributions (50:50) to evaluate *BI* and *BIC diff* performance to identify bimodal distributions. Shown is an example of a randomly generated set of 100 benchmarking data sets with 10,000 samples obtained for unimodal (white background) or bimodal (green background) distributions. The red and blue lines indicate the corresponding unimodal and bimodal fits, respectively.

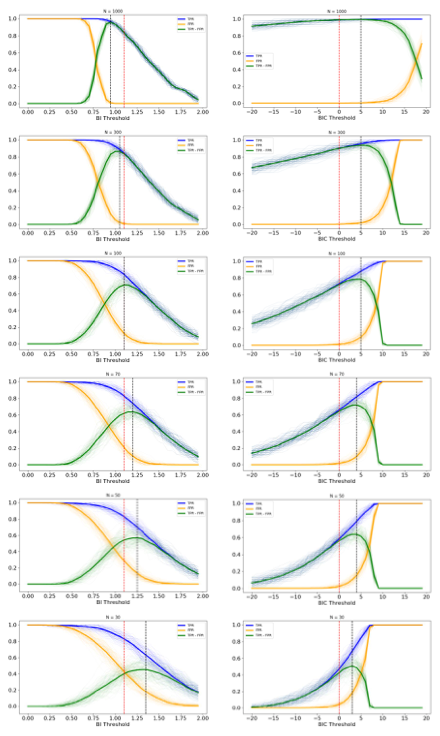

**Supplementary Figure S3.** Performance of *BI* and *BIC diff* in identifying bimodal distributions. Shown are the true-positive rate [TPR], the false-positive rate [FPR,] and their differences [TPR – FPR]) in 100 different experiments comprising different sample sizes (*N* = 1000, 300, 100, 70, 50, 30). Solid lines indicate the average of the 100 experiments. Vertical dotted lines correspond to the theoretical threshold (red) and observed best threshold (black): max(TPR -FPR). x-axis: *BI*/*BIC diff* threshold; y-axis: TRP/FPR/(TRP – FPR).

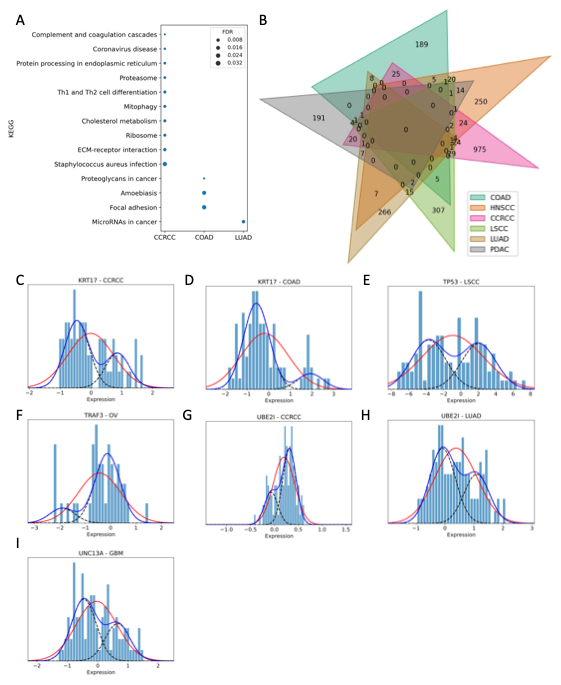

**Supplementary Figure S4.** (A) KEGG pathway enrichment analysis using indication-specific TABPs. The sizes of the colored circles represent FDR. (B) Venn plot of tumor-specific bimodal proteins in 6 indications. (C–I) Distributions of five bimodal proteins (KRT17, TP53, TRAF3, UBE2I, UNC13A) in the CPTAC pan-cancer data set. The red and blue lines indicate the corresponding unimodal and bimodal fits, respectively.

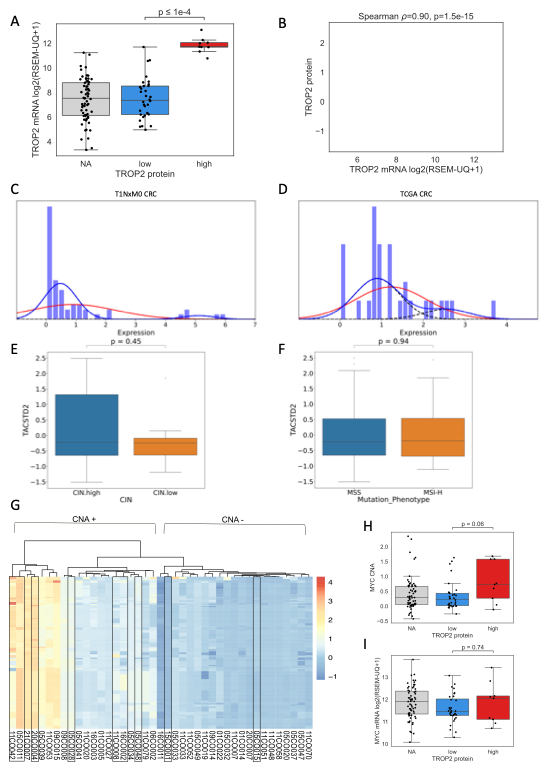

**Supplementary Figure S5.** TROP2 bimodality in the CPTAC COAD cohort. (A) TROP2 mRNA was significantly higher in the TROP2-high samples identified by the bimodality modal (*P* = 1e-4). (B) Correlation analysis between TROP2 mRNA and TROP2 protein in the CPTAC COAD cohort (Spearman correlation, *P* = 0.90, *P* = 1.5e-15). (C–D) Bimodal TROP2 protein expression observed in T1NxM0 CRC and TCGA CRC proteomics data. The red and blue lines indicate the corresponding unimodal and bimodal fits, respectively. (E) TROP2 protein expression in samples classified as CIN high versus CIN low (*P* = 0.45). (F) TROP2 protein expression in samples stratified according to microsatellite instability status (*P* = 0.94). (G) Heat map showing top 323 copy number altered genes associated with TROP2 abundance using MOFA (requiring feature weights of >0.9 for top factor with high weights for TROP2 protein and mRNA abundance). (H) *MYC* gene copy number alteration values in TROP2-high, -low, and not-quantified samples (*P* = 0.06 TROP2 high vs. TROP2 low). (I) *MYC* gene mRNA abundance in TROP2-high, -low, and not-quantified samples (*P* = 0.74, TROP2 high vs. TROP2 low).

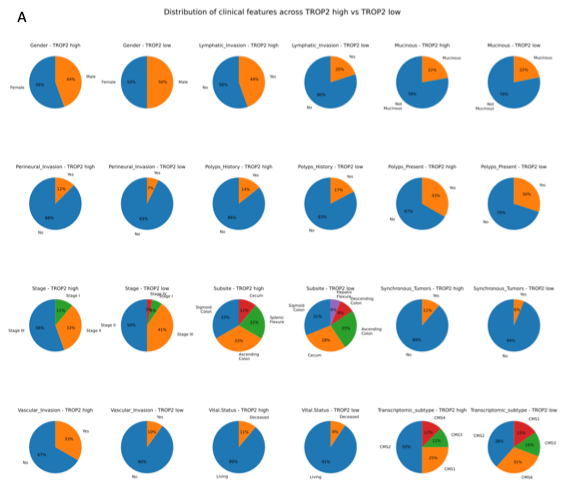

**Supplementary Figure S6.** TROP2 bimodality in the COAD cohort, showing the distribution of demographic and clinical features of TROP2-high and TROP2-low samples including vital status, vascular invasion, synchronous tumors, tissue subsite, tumor stage, presence of polyps, polyp history, perineural invasion, mucinous phenotype, lymphatic invasion, and gender (all *P* > 0.05, chi-square test).

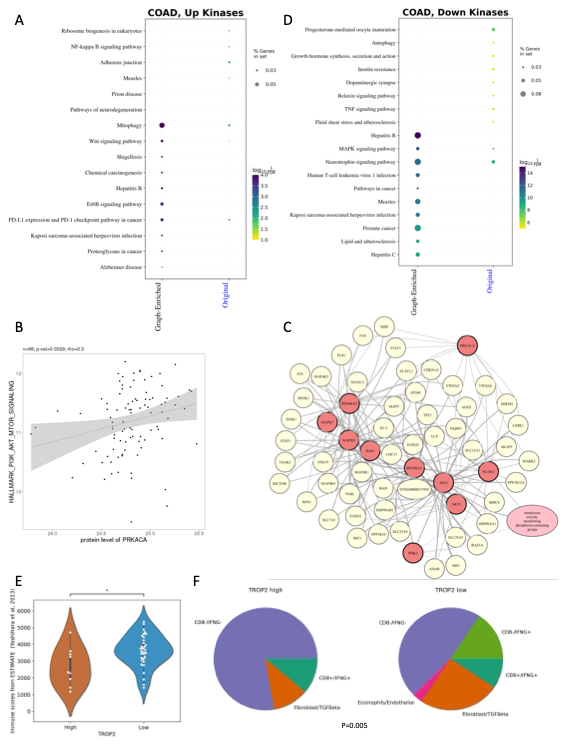
**Supplementary Figure S7.** (A) Pathway enrichment analysis using the Biological Insight Knowledge Graph (BIKG)–enriched proteins and original upregulated kinases. The sizes of the colored circles represent the percentage of the gene covered in the gene set; colors represent –log10(FDR). (B) Correlation between protein abundance of PRKACA and hallmark signature scores of PI3K_ATK_MTOR_SIGNALING in COAD (*r* = 0.3, *P* = 0.0029). (C) Network extracted from BIKG with downregulated kinases as seeds. Red nodes indicate the downregulated kinases, yellow, closely connected proteins to the downregulated kinases; pink, biological processes. (D) Pathway enrichment analysis using the BIKG-enriched proteins and original downregulated kinases. (E) Immune scores from ESTIMATE in TROP2-high and -low samples. *P* values were derived from the *t* test with Bonferroni correction. *****P* < 0.0001. For box plots, the centerline indicates the median, box limits indicate upper and lower quartiles, and whiskers indicate the 1.5 interquartile range. (F) Proportion of samples featuring each immune subtype from Petralia et al.^1^ in TROP2-high versus TROP2-low samples (chi-square test, *P* = 0.005).

**Reference**

1	Petralia, F. *et al.* Pan-cancer proteogenomics characterization of tumor immunity. *Cell* **187**, 1255-1277 (2024).
